# Exploring the remarkable diversity of *Escherichia coli* Phages in the Danish Wastewater Environment, Including 91 Novel Phage Species

**DOI:** 10.1101/2020.01.19.911818

**Authors:** Nikoline S. Olsen, Witold Kot, Laura M. F. Junco, Lars H. Hansen

**Affiliations:** Department of Environmental Science, Aarhus University, Frederiksborgvej 399, Roskilde, Denmark; Department of Plant and Environmental Sciences, University of Copenhagen, Thorvaldsensvej 40, 1871 Frederiksberg C, Denmark

**Keywords:** bacteriophage, wastewater, *Escherichia coli*, diversity, genomics, taxonomy, coliphage

## Abstract

Phages drive bacterial diversity - profoundly influencing diverse microbial communities, from microbiomes to the drivers of global biogeochemical cycling. The vast genomic diversity of phages is gradually being uncovered as >8000 phage genomes have now been sequenced. Aiming to broaden our understanding of *Escherichia coli* (MG1655, K-12) phages, we screened 188 Danish wastewater samples (0.5 ml) and identified 136 phages of which 104 are unique phage species and 91 represent novel species, including several novel lineages. These phages are estimated to represent roughly a third of the true diversity of *Escherichia* phages in Danish wastewater. The novel phages are remarkably diverse and represent four different families *Myoviridae, Siphoviridae, Podoviridae* and *Microviridae*. They group into 14 distinct clusters and nine singletons without any substantial similarity to other phages in the dataset. Their genomes vary drastically in length from merely 5 342 bp to 170 817 kb, with an impressive span of GC contents ranging from 35.3% to 60.0%. Hence, even for a model host bacterium, in the go-to source for phages, substantial diversity remains to be uncovered. These results expand and underlines the range of *Escherichia* phage diversity and demonstrate how far we are from fully disclosing phage diversity and ecology.

## 1. Introduction

Phages are important ecological contributors, they renew organic matter supplies in nutrient cycles and drive bacterial diversity by enabling co-existence of competing bacteria by “*Killing the winner*” and by serving as genomic reservoirs and transport units [1,2]. Phage genomes are known to entail auxiliary metabolism genes (AMGs), toxins, virulence factors and even antibiotic resistance genes [3–7] and through lysogeny and transduction they can transfer metabolic traits to their hosts and even confer immunity against homologous phages [1]. Still, in spite of their ecological role, potential as antimicrobials and the fact that they carry a multitude of unknown genes with great potential for biotechnological applications, phages are vastly understudied. Less than 9000 phage genomes have now been published, and though the number increases rapidly, we may have merely scratched the surface of the expected diversity [8]. It has been estimated that at least a billion bacterial species exist [9], hence only phages targeting a tiny fraction of potential hosts have been reported. Efforts to disclose the range and diversity of phages targeting a single host, have revealed a stunning display of diversity. The most scrutinized phage host is the *Mycobacterium smegmatis*, for which the Science Education Alliance Phage Hunters program has isolated more than 4700 phages and fully sequenced 680 phages which represent 30 distinct phage clusters [10,11]. This endeavour has provided a unique insight into viral and host diversity, evolution and genetics [12–15]. No other phage host has been equally targeted, but *Escherichia coli* phages have been isolated in fairly high numbers. The International Committee on Taxonomy of Viruses (ICTV) has currently recognised 158 phage species originally isolated on *E. coli* [16], while 2732 *Escherichia* phage genomes have been deposited in GenBank [17]. As phages are expected to have an evolutionary potential to migrate across microbial populations, host species may not be an ideal indicator of relatedness, but it serves as an excellent starting point to explore phage diversity. Hierarchical classification of phages is complicated by the high degree of horizontal gene transfer, consequently several classification systems have been proposed [18–20], and we may not yet have reached a point where it is reasonable to establish the criteria for a universal system [20]. Nonetheless, a system enabling a mutual understanding and exchange of knowledge is needed. Accordingly, we have in this study chosen to classify our novel phages according to the ICTV guidelines [21].

Here we aim to expand our understanding of *Escherichia* phage diversity. Earlier studies are few, have relied on more laborious and time-consuming methods, or are based on *in silico* analyses of already published phage genomes [8]. Accordingly, Korf *et al*., (2019) isolated 50 phages on 29 individual *E. coli* strains, while Jurczak-Kurek *et al*., (2016) isolated 60 phages on a single *E. coli* strain, both finding a broad diversity of *Caudovirales* and also representatives of novel phage lineages [22,23]. We hypothesized, that a high-throughput screening of nearly 200 samples in time-series, using a single host, would expand the number of known phages by facilitating the interception of the prevailing phage(s) of the given day.

## 2. Materials and Methods

The screening for *Escherichia* phages was performed with the microplate based *High-throughput screening method* as described in (not published). With the exception that lysates of wells giving rise to plaques were sequenced without further purification. In short, an overnight enrichment was performed in microplates with host culture, media and wastewater (0.5 ml per well), followed by filtration (0.45 µm), a purification step by re-inoculation, a second overnight incubation and filtration (0.45µm), and then a spot-test (soft-agar overlay) to indicate positive wells. All procedures were performed under sterile conditions.

### 2.1.1 Samples Bacteria and media

Inlet Wastewater samples (188) were collected (40-50 ml) in time-series (2-4 days spanning 1-3 weeks), from 48 Danish wastewater treatment facilities (rural and urban) geographically distributed on Zealand, Funen and in Jutland, during July and August 2017. The samples were centrifuged (9000 x g, 4 °C, 10 min) and the supernatant filtered (0.45 µm) before storage in aliquots (−20°C) until screening. The host bacterium is *E. coli* (MG1655, K-12), and the media Lysogeny Broth (LB), amended with CaCl_2_ and MgCl_2_ (final concentration 50 mM). Collected phages were stored in SM-buffer [24] at 4°C.

### 2.1.2 Sequencing and genomic characterisation

DNA extractions, clean-up (ZR-96 Clean and Concentrator kit, Zymo Research, Irvine, CA USA) and sequencing libraries (Nextera® XT DNA kit, Illumina, San Diego, CA USA) were performed according to manufacturer’s protocol with minor modifications as described in Kot et al., (2014) [25]. The libraries were sequenced as paired-end reads on an Illumina NextSeq platform with the Mid Output Kit v2 (300 cycles). The obtained reads were trimmed and assembled in CLC Genomics Workbench version 10.1.1. (CLC BIO, Aarhus, Denmark). Overlapping reads were merged with the following settings: mismatch cost: 2, minimum score: 15, gap cost: 3 and maximum unaligned end mismatches: 0, and then assembled *de novo*. Additional control assemblies were constructed using SPAdes version 3.12.0 [26]. Phage genomes were defined as contigs with an average coverage > x 20 and a sequence length ≥ 90% of closest relative. Gene prediction and annotation were performed using a customized RASTtk version 2.0 [27] workflow with GeneMark [28], with manual curation and verification using BLASTP [29], HHpred [30] and Pfam version 32.0 [31], or de novo annotated using VIGA version 0.11.0 [32] based on DIAMOND searches (RefSeq Viral protein database) and HMMer searches (pVOG HMM database). All genomes were assessed for antibiotic resistance genes, bacterial virulence genes, type I, II, III and IV restriction modification (RM) genes and auxiliary metabolism genes (AMGs) using ResFinder 3.1 [33,34], VirulenceFinder 2.0 [35], Restriction-ModificationFinder 1.1 (REBASE) [36] and VIBRANT version 1.0.1 [37], respectively. The 104 unique phage genomes were aligned to viromes of BioProject PRJNA545408 in CLC Genomics Workbench and deposited in Genbank [17].

### 2.1.3 Genetic analyses

Nucleotide (NT) and amino acid (AA) similarities were calculated using tools recommended by the ICTV [38], i.e. BLAST [29] for identification of closest relative (BLASTn when possible, discontinuous megaBLAST (word size 16) for larger genomes) and Gegenees version 2.2.1 [39] for assessing phylogenetic distances of multiple genomes, for both NTs (BLASTn algorithm) and AAs (tBLASTx algorithm) a fragment size of 200 bp and step size 100 bp was applied. NT similarity was determined as percentage query cover multiplied by percentage NT identity. Novel phages are categorised according to ICTV taxonomy. The criterion of 95% DNA sequence similarity for demarcation of species was applied to identify novel species representatives and to determine uniqueness within the dataset. Evolutionary analyses for phylogenomic trees were conducted in MEGA7 version 2.1 (default settings) [40]. These were based on the large terminase subunit (*Caudovirales*), a gene commonly applied for phylogenetic analysis [41,42] and on the DNA replication gene (*gpA*) (*Microviridae*). The NT sequences were aligned by MUSCLE [43] and the evolutionary history inferred by the Maximum Likelihood method based on the Tamura-Nei model [44]. The trees with the highest log likelihood are shown. Pairwise whole genome comparisons were performed with Easyfig 2.2.2 [45] (BLASTn algorithm), these were curated by adding color-codes and identifiers in Inkscape version 0.92.2. The R package iNEXT [46,47] in R studio version 1.1.456 [48] was used for rarefaction, species diversity (q = 0, datatype: incidence_raw), extrapolation hereof (estimadeD) and estimation of sample coverage. The visualisation of genome sizes and GC contents was prepared in Excel version 16.31.

**Figure 1.**
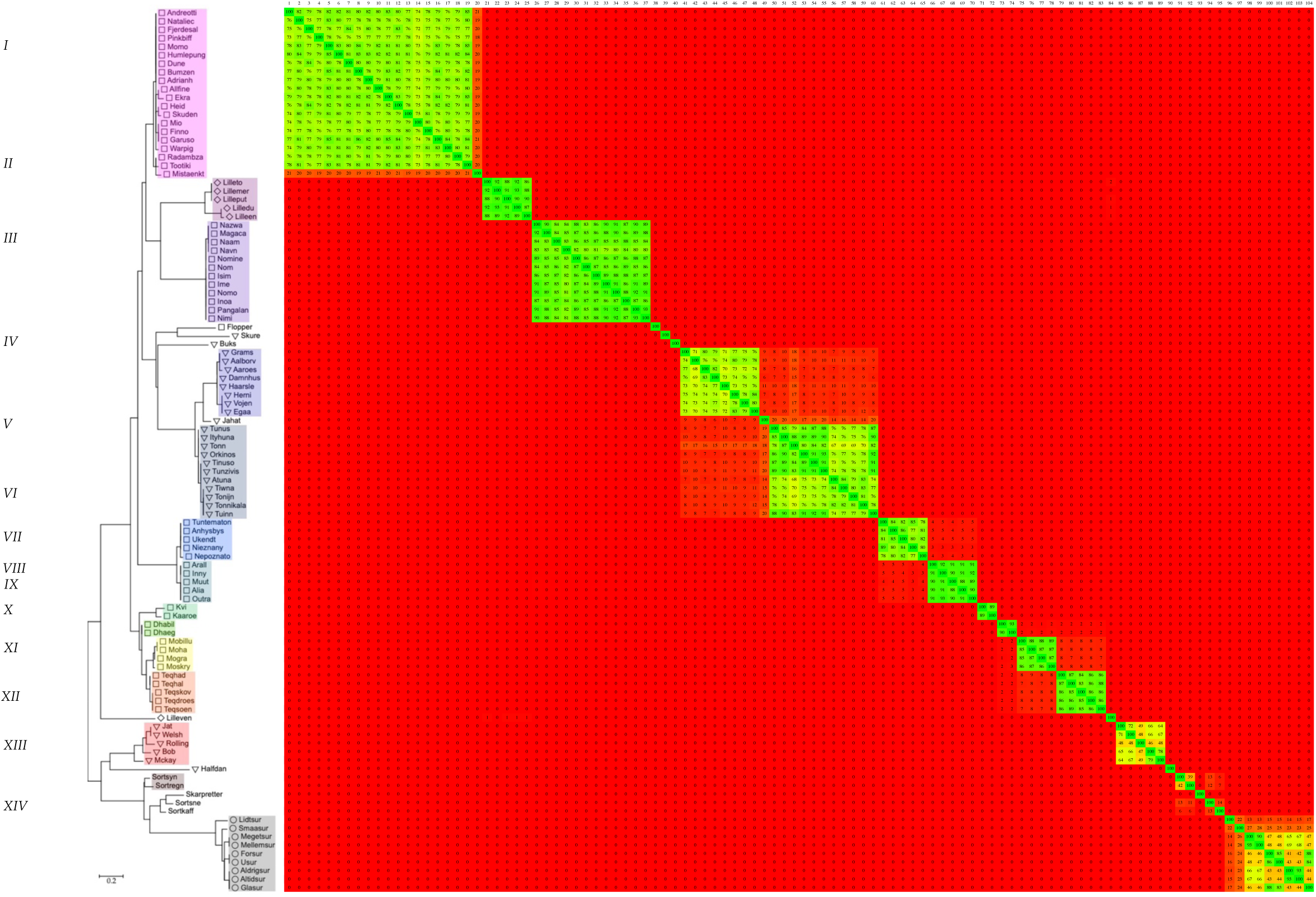
Phylogenetic tree (Maximum log Likelihood: −1325.01, large terminase subunit (*Caudovirales*) or DNA replication protein *gpA* (*Microviridae*)), scalebar: substitutions per site, morphology is indicated by symbols (*Myoviridae:* □, *Siphoviridae:* ▽, *Podoviridae:* ○, *Microviridae:* ◊) and phylogenomic nucleotide distances (Gegenees, BLASTn: fragment size: 200, step size: 100, threshold: 0%)

**Figure 2.**
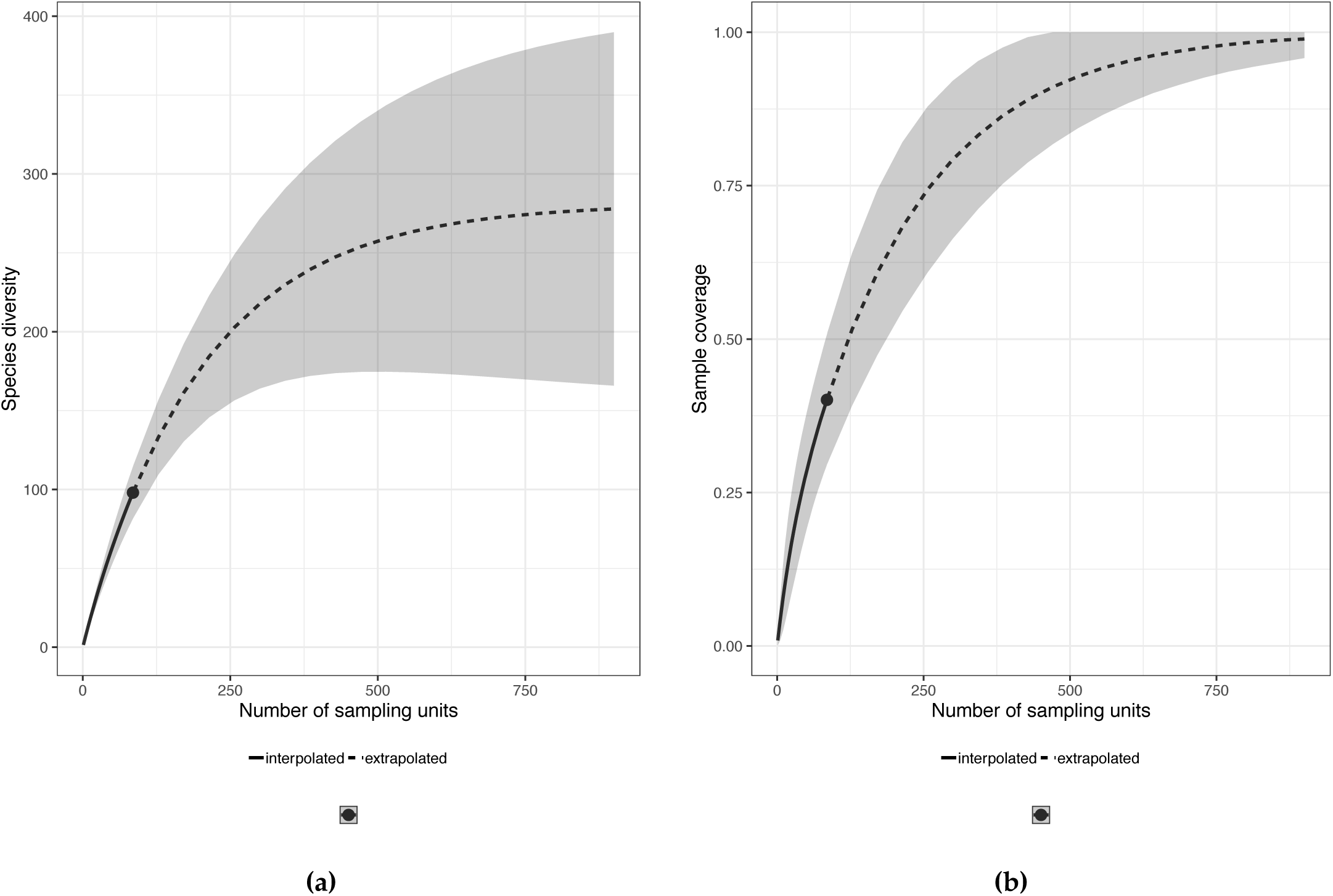
**(a)** Sample-size-based rarefaction and extrapolation curve with confidence intervals (0.95). **(b)** Sample completeness curve with confidence intervals (0.95).

## 3. Results and Discussion

### 3.2. Phylogenetics, taxonomy and species richness

The sequenced phages were analysed strictly *in silico*, focusing on their relatedness to known phages, their taxonomy and any distinctive characteristics. Based on the confirmed morphology of closely related phages, at least four different families are represented, *Myoviridae* (58%), *Siphoviridae* (25%), *Podoviridae* (7.4%) and even the single stranded DNA (ssDNA) *Microviridae* (5.9%) (Table 1). Five (3.7%) of the phages are so distinct that they could not with certainty be assigned to any family. A similar distribution was found by Korf *et al*., (2019), i.e. 70% *Myoviridae*, 22% *Siphoviridae* and 8% *Podoviridae*, just as an analysis of the genomes of *Caudovirales* infecting *Enterobacteriaceae*, by Grose & Casjens (2014), also identified more clusters belonging to the *Myoviridae*, than the *Siphoviridae* and fewest of the *Podoviridae* [8,22]. Jurczak-Kurek *et al*., (2016) found more siphoviruses than myovirus, but also found the *Podoviridae* to be the least abundant [23]. Nonetheless, these distributions are just as likely to be caused by culture or isolation methods, as they are to reflect true abundances. Based on DNA homology and phylogenetic analysis, the 104 unique phages identified in this study group into 14 distinct clusters and nine singletons, having a <1% Gegenees score between clusters and singletons, with the exceptions of *cluster IV, V* and Jahat which have inter-Gegenees scores of up to 20%, *cluster IX, X* and *XI* which have inter-Gegenees scores of 1-8% and *cluster XIII*, Sortsne and Sortkaff which have inter-Gegenees scores of 6-14% (Figure 1, Table 2). Excluding one phage in *cluster I* and two phages in *cluster XIV*, the intra-Gegenees score of all clusters is above 39% (Figure 1).

**Table 1.**
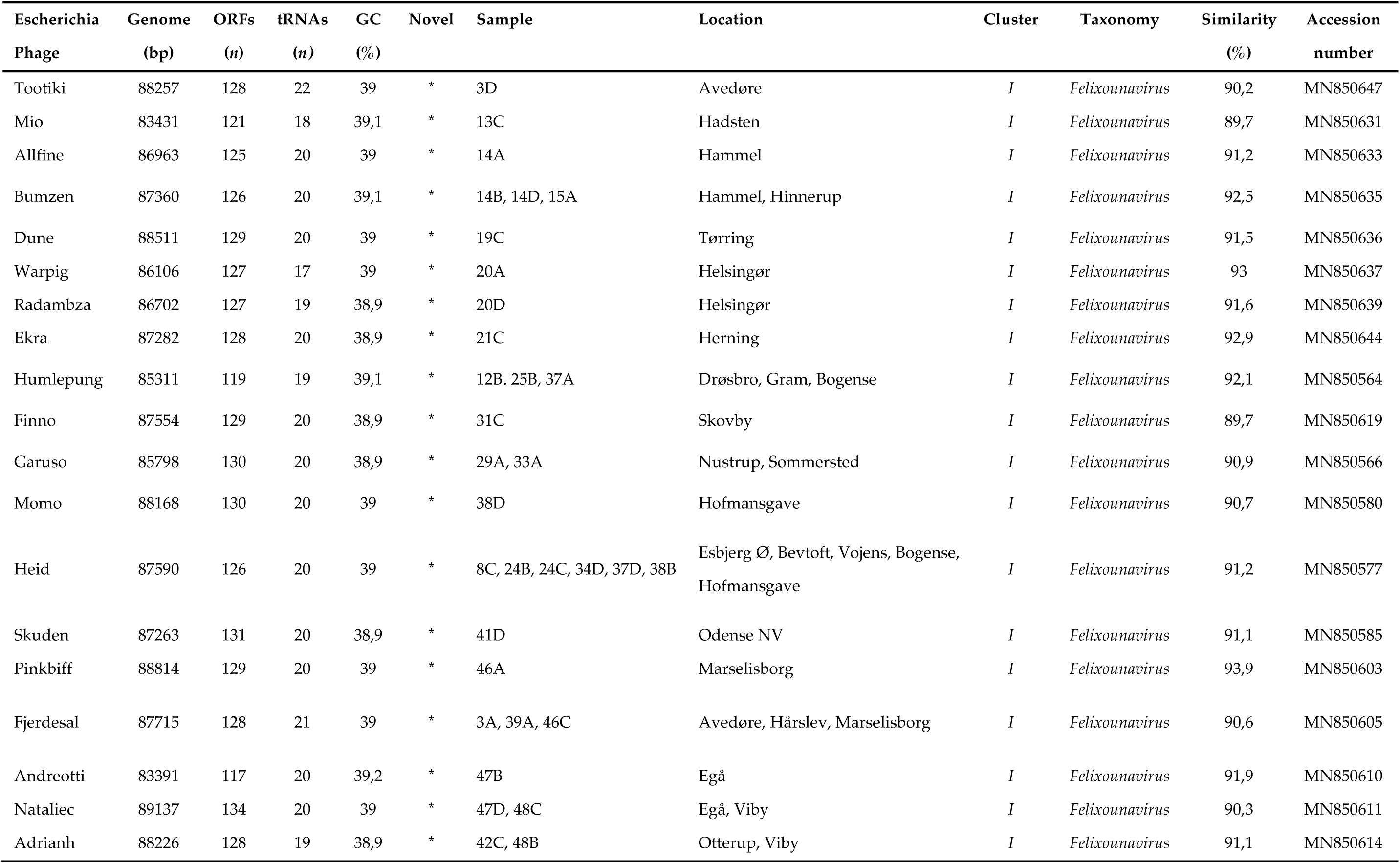
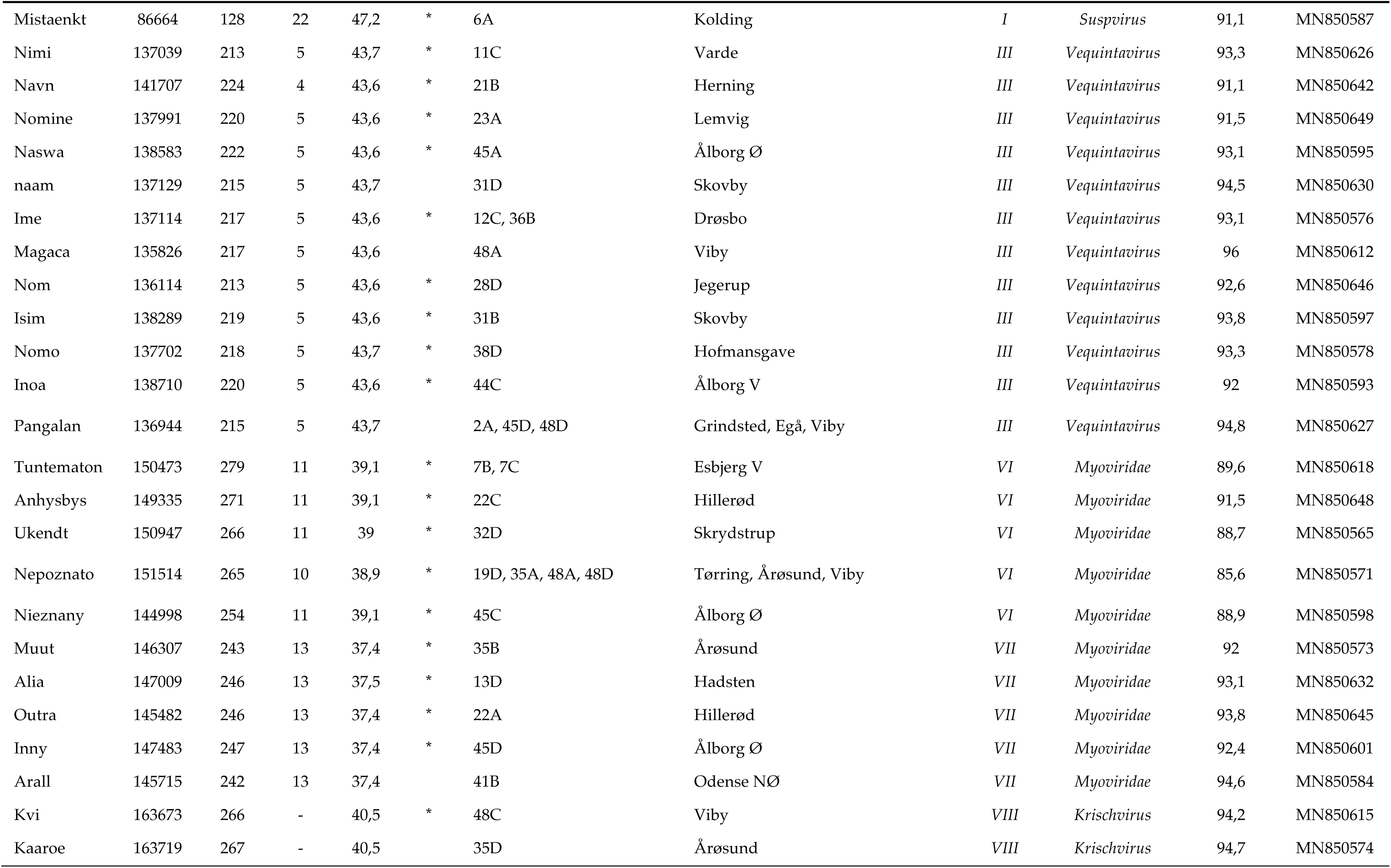
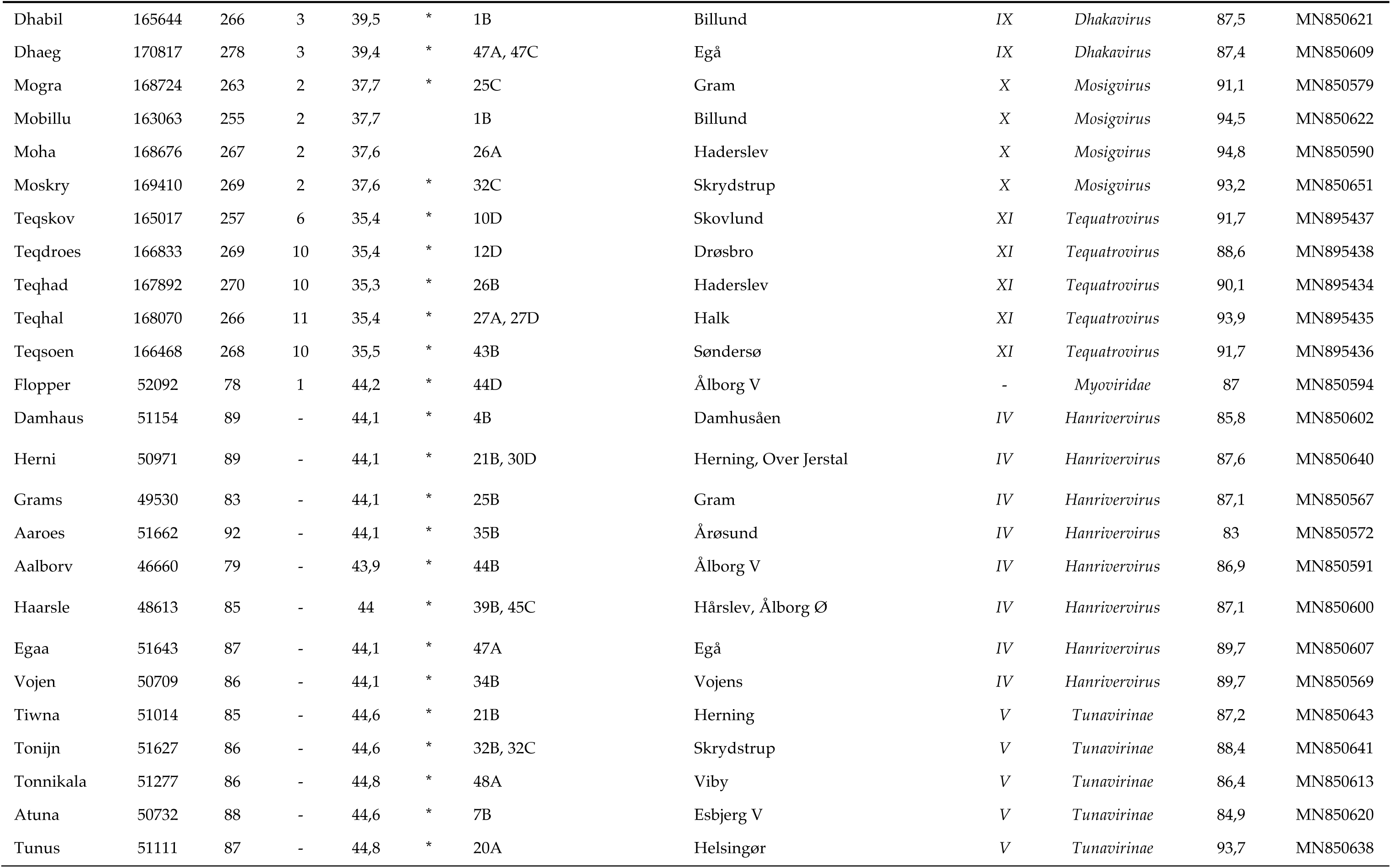
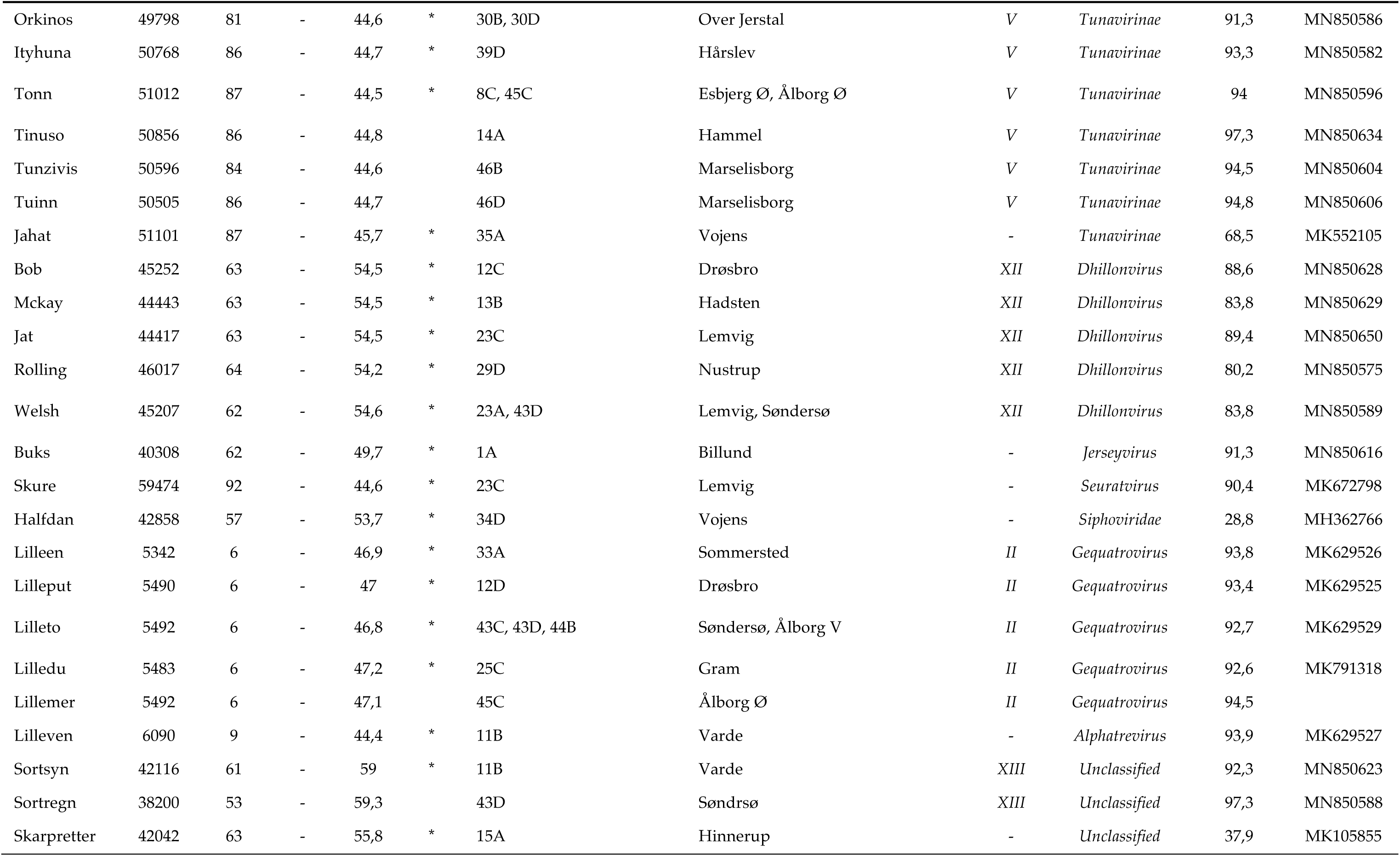
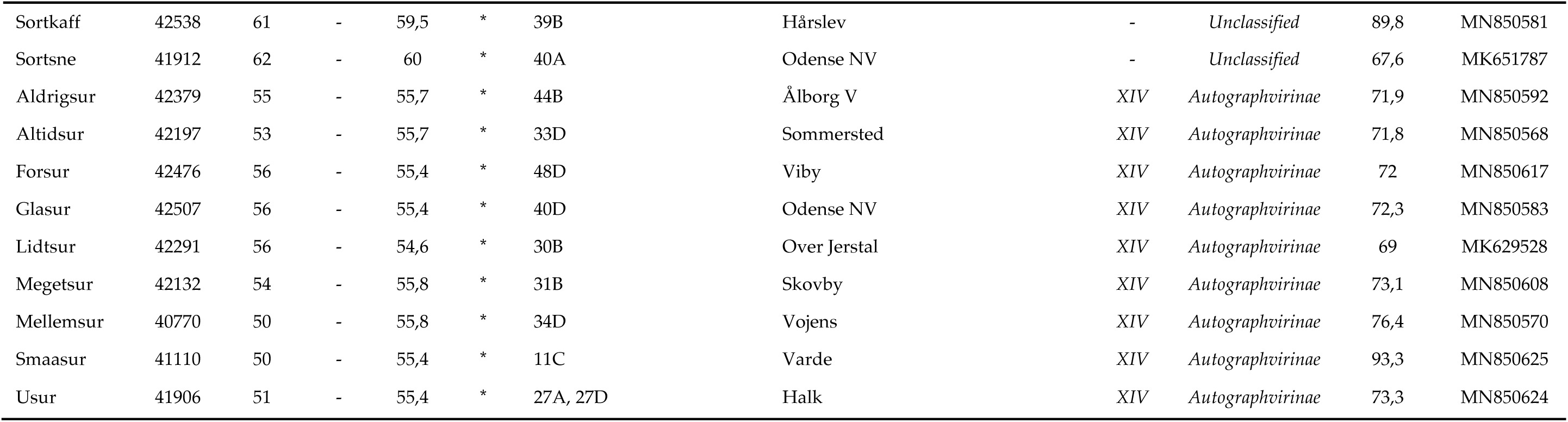
List of 104 unique *Escherichia* phages identified in 94 Danish wastewater samples. Novelty (*) is based on the 95% demarcation of species. Similarity is sequence identity (%) times sequence coverage (%) to closest related (Blastn). Taxonomy is based on similarity (BLASTn) to closest related and the ICTV Master Species list.

**Table 2.**
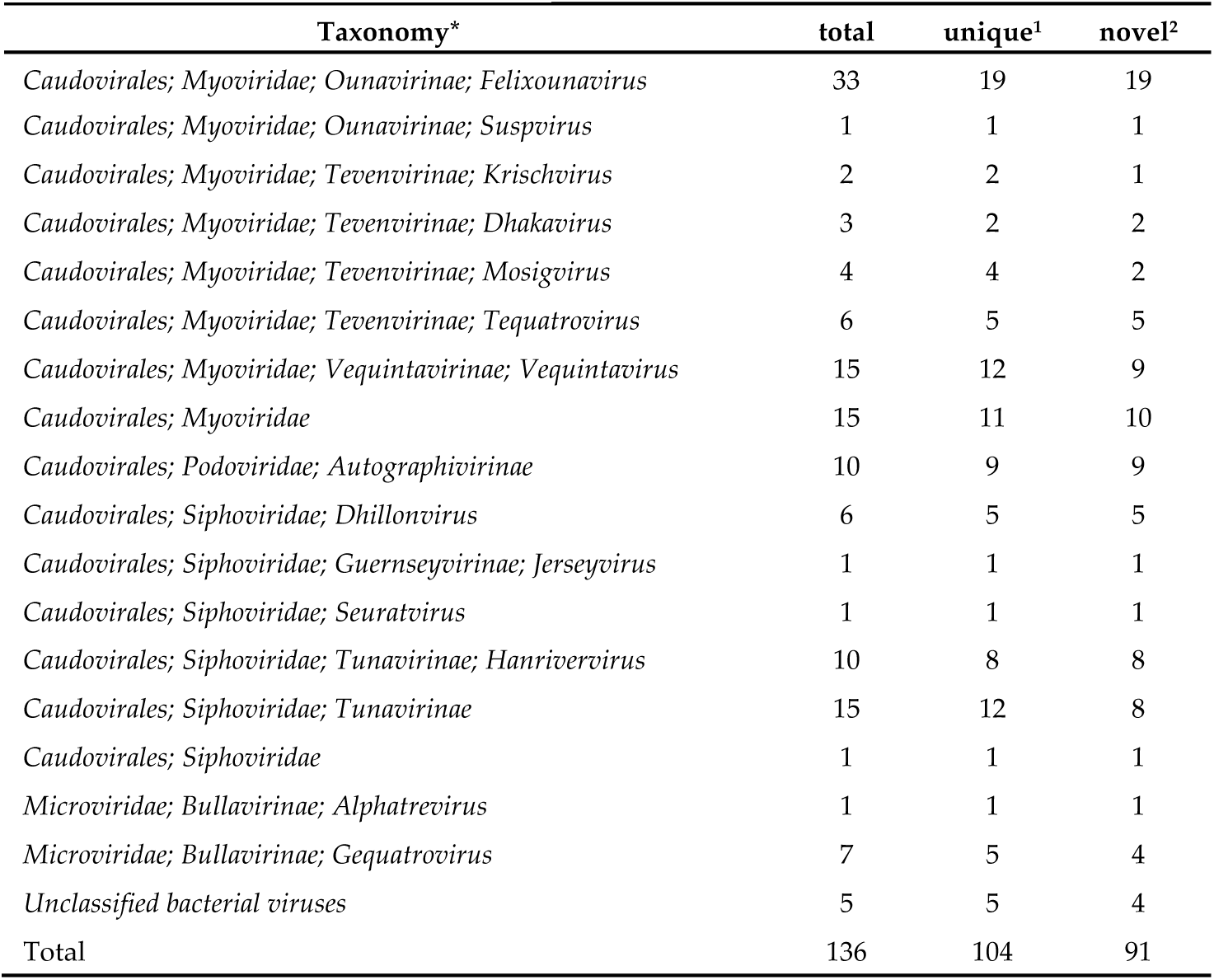
Taxonomic distribution of phages identified in 94 Danish wastewater samples, based on similarity to closest related and the ICTV Master Species list. 1. ≤95% similarity to other phages in the dataset. 2. ≤95% similarity to other phages in the dataset and to published phage genomes.

The high-throughput screening method favours easily culturable plaque-forming lytic phages. Still, we identified 104 unique Escherichia phages of which only 16% were ≥95% similar (BLAST) to already published phages (Table 2). Phages were identified in wastewater samples from 43 of the 48 investigated treatment facilities. From the majority of positive samples (58) a single phage was sequenced, however in some samples the lysate held more than one phage. Twenty-five of the lysates held two phages, eight lysates held three phages and one had as many as four phages. Of the 104 unique phages, 91 represent novel species (Table 1, 2). Of these, 51 differed by ≥10% from published phage genomes and some have NT similarities as low as 29% (Table 2).

These newly sequenced phages represent a substantial quota of divergent lytic *Escherichia* phages in Danish wastewater, but are still far from disclosing the true diversity hereof (Figure 2). An extrapolation of species richness (q = 0) predicts a total of 292 distinct species (requiring a sample size of ∼900 phages). The relatively small sample-size in this study (n = 136), may subject the estimation to a large prediction bias. The sampling-method also introduces a bias by selecting for abundance and burst size, thereby potentially underestimating diversity. Nonetheless, the results provide an indication of the minimal diversity of lytic *Escherichia* (MG1655, K-12) phages in Danish wastewater, estimated to be as a minimum be in the range of 160 to 420 unique phage species (Figure 2b). The novelty and diversity of these wastewater phages is truly remarkable and verifies our hypothesis, as well as the efficiency of the *High-throughput Screening Method* for exploring diverse phages of a single host (not published).

### 3.2. Phage genome characteristics

The sequenced phage genomes range in sizes from 5 342 bp of the *Microviridae* lilleen to a span in the *Caudovirales* from 38 200 bp of the unclassified sortregn to 170 817 bp of the *Dhakavirus* dhaeg (Figure 3, Table 2). GC contents also vary greatly, from only 35.3% (*Tequatrovirus* teqhad) and up to 60.0% (the unclassified sortsne) (Figure 3,Table 2), heavily diverging from the host GC content of 50.79%. Phages often have a lower GC contents than their host [49], as observed for the majority (81%) of the wastewater phages (Table 2). A lower GC content of phage genomes tends to correlate with an increase in genome size [50], as is also the case for the *Caudovirales* in this study (Table 2). Rocha & Danchin (2002) hypothesized that phages, along with plasmids and insertion sequences, can be considered intracellular pathogens, and like host dependent bacterial pathogens and symbionts, they experience competition for the energetically expensive and less abundant GTP and CTP and as a consequence develop genomes with comparatively higher AT contents [49]. However, the differences in AT richness reported by Rocha & Danchin (2002) for dsDNA and isometric ssDNA phages were merely 4.2% and 5% [49], while the dsDNA and isometric ssDNA phages in this study deviate from their isolation host by having up to 31% higher and up to 19% lower AT content (Table 2). Accordingly, the considerable differences in GC/AT contents may instead primarily be a reflection of past host relations [14]. The sequencing of lysates and not only plaquing phages, may have contributed in enabling the capture of such a broad GC-content and overall diversity.

**Figure 3.**
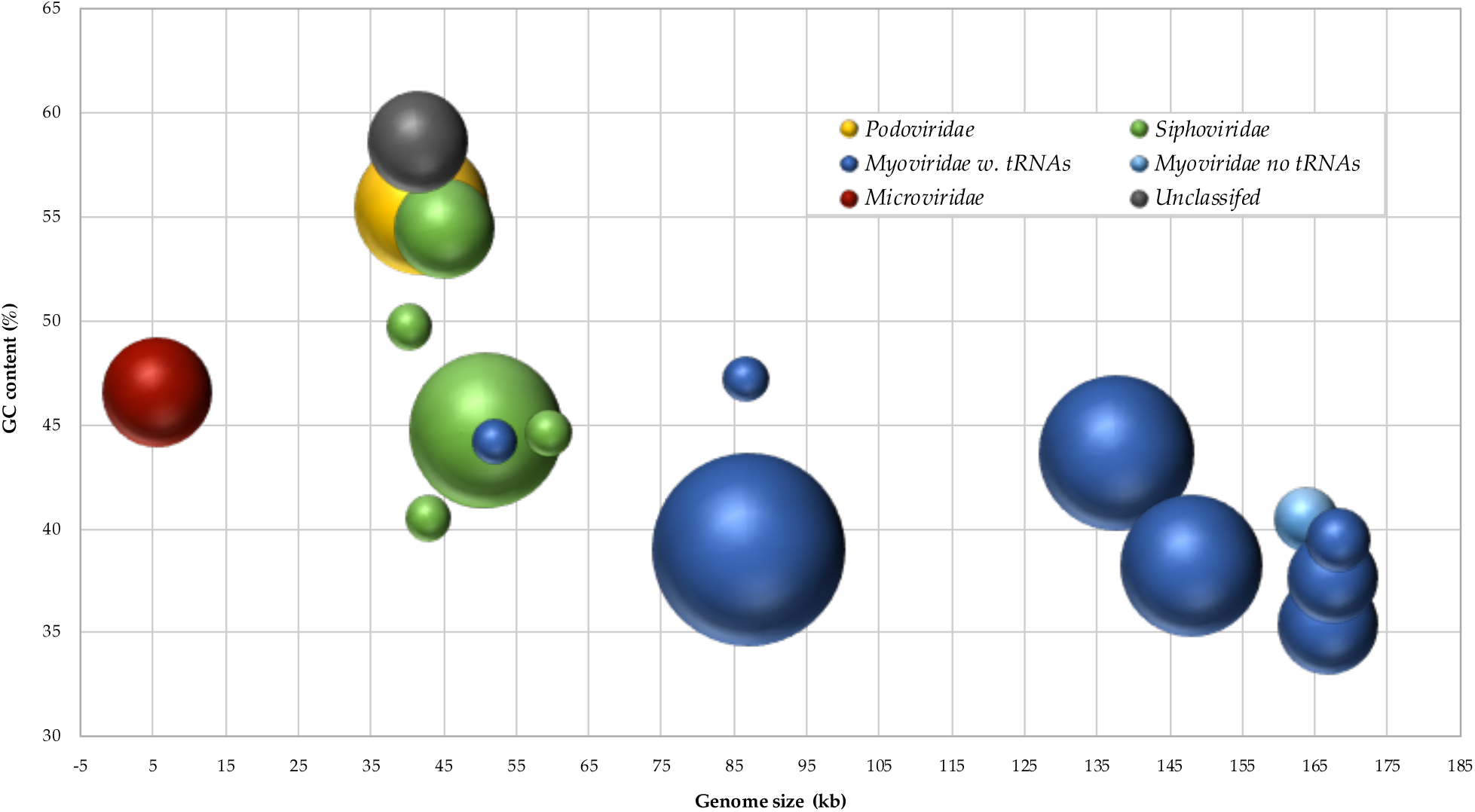
Bubble-diagram of the 104 unique *Escherichia* phages displaying genome size- and GC-content distribution. Area of bubbles indicates number of phages. The yellow bubble is *Podoviridae* phages, green ones are *Siphoviridae* phage clusters, dark-blue ones are *Myoviridae* phage clusters with tRNAs, the light-blue bubble is *Myovridae* phages without tRNAs, the red bubble is *Microviridae* phages and the grey one is unclassified phages.

The genome screening algorithms identified no sequences coding for homologs of known virulence or antibiotic resistance genes. Though not a definitive exclusion, this interprets as a reduced risk of presence, a preferable trait for phage therapy application. Currently available tools for AMG screening of viromes did not provide a comprehensive and exclusive assessment of the AMG pool in the dataset. The majority of genes identified are not AMGs, but code for phage DNA modification pathways (Table S1). The function of some of the suggested AMGs is unknown, these include a nicotinamide phosphoribosyltransferase (NAMPT) present in *cluster I* (*Felixounavirus* and *Suspvirus*) and *VII* (unclassified *Myoviridae*), a 3-deoxy-7-phosphoheptulonate (DAHP) synthase present in *cluster X* (*Mosigvirus)* and a complete dTDP-rhamnose biosynthesis pathway present in *cluster VI* and *VII* (unclassified *Myoviridae*). The presence of a dTDP-rhamnose biosynthesis pathway in the DNA metabolism region of phage genomes is peculiar, one possible explanation is, that these phages utilize rhamnose for glycosylation of hydroxy-methylated nucleotides in the same manner as the T4 generated glucosyl-hmC [51]. Dihydrofolate reductase, identified in six of the *Myoviridae clusters*, is involved in thymine nucleotide biosynthesis, but is also a structural component in the tail baseplate of T-even phages [52]. Multiple verified DNA modification genes were identified. Methyltransferases, some putative, were detected in all *Myoviridae of cluster III* (*vequintavirus*), *VI* (unclassified), *VII* (unclassified), *VIII* (*Krischvirus*), in all *Siphoviridae* of *cluster IV* (*Hanrivervirus*) and in the unclassified phages of *cluster XIII*. Indeed, *cluster XIII* and the singletons skarpretter, sortsne, and sortkaff code for both DNA N-6-adenine-methyltransferases (*dam)* and DNA cytosine methyltransferases (*dcm*). Finally, two novel epigenetic DNA hypermodification pathways were identified. The mosigviruses of *cluster X* code for arabinosylation of hmC [51], while the novel seuratvirus Skure codes for the recently verified complex 7-deazaguanine DNA hypermodification system [53,54].

### 3.3. Forty-eight novel Myoviridae phages species

The *Myoviridae* genomes (56 unique, 49 novel) represents the greatest span in genome sizes in this study, from the unclassified flopper of 52092 bp to the dhakavirus dhaeg of 170817 bp), all, except the krischviruses, code for tRNAs (Figure 3, Table 2). *The Myoviridae* group into nine distinct clusters and four singletons, representing at least three subfamilies; *Tevenvirinae, Vequintavirinae* and *Ounavirinae* in addition to 11 unclassified *Myoviridae* (*cluster VI & VII*) (Figure 1, Table 2). The eleven novel *Tevenvirinae* distributes into four distinct genera, two of the *Krischvirus* (*cluster VIII*, 164kb, 40.5% GC, no tRNAs), two of the *Dhakavirus* (*cluster IX*, 166-170 kb, 39.4-39.5% GC, 3 tRNAs), two of the *Mosigvirus* (*cluster X*, 169 kb, 37.6-37.7% GC, 2 tRNAs) and five of the *Tequatrovirus* (*cluster XI*, 165-168 kb, 35.3-35.5% GC, 6-11 tRNAs), while the nine novel *Vequintavirinae* are all vequintaviruses (*cluster III*, 136-142 kb, 43.6-43.7% GC, 5 tRNAs) closely related (91.1-93.8%, BLAST) to classified species. The vequintaviruses were identified in samples from 12 treatment facilities. Only reads from two samples of a dataset of human gut viromes based on timeseries of faecal samples from 10 healthy persons [55] mapped to the wastewater phages, and only to vequintaviruses. These reads covered 43-54% and 13-26% of all the *Vequintavirus* genomes except pangalan and navn, respectively. This suggests that the vequintaviruses are related to entero-phages. All but one of the *Ounavirinae* (*cluster I*, 82-89 kb, 38.9-39.2% GC, 17-22 tRNAs) are felixounaviruses (89.7-93.9%, BLAST) with an intra-Gegenees score of 71-86% (Figure 1). The *Felixounavirus* is a relatively large genus with 17 recognized species. In this study, felixounaviruses were identified in samples from 23 of the 48 facilities, indicating that they are both ubiquitous in Danish wastewater, numerous and easily cultivated. The last ounavirus, mistaenkt (86.7 kb, 47.2% GC, 22 tRNAs) is a *Suspvirus*. The five *cluster VI* phages (144.9-151.5 kb, 39±0.1% GC, 10-11 tRNAs) belong to a phylogenetic distinct clade, the recently proposed ‘*Phapecoctavirus*’, and have substantial similarity (86-90%, BLAST) with the anticipated type species Escherichia phage phAPEC8 (JX561091) [22,56]. *Cluster VII* have significantly (*p* = 0.038) larger genomes (145.8-147.5 kb) with marginally, though not significantly (*p* = 0.786), lower GC contents of 37.4-37.5%, all have 13 tRNAs (Table 2) and code for NAMPT not present in *cluster VI* (Table S1). As a group, *cluster VII* are even more homogeneous than *cluster VI* and all are closely related (92-95%, BLAST) to the same five unclassified *Enterobacteriaceae* phages vB_Ecom_PHB05 (MF805809), vB_vPM_PD06 (MH816848), ECGD1 (KU522583), phi92 (NC_023693) and vB_vPM_PD114 (MH675927) [57,58], with whom they represent a novel unclassified genera. The distinctive and novel singleton flopper only shares NT similarity (36-87%, BLAST) with six other phages, the *Escherichia* phages ST32 (MF04458) [59] and phiEcoM_GJ1 (EF460875) [60], the *Erwinia* phages Faunus (MH191398) [61] and vB_EamM-Y2 (NC_019504) [62], and the *Pectobacterium* phages PM1 (NC_023865) [63] and PP101 (KY087898). This group of unclassified phages all have genomes of 52.1-56.6 kb with 43.6-44.9% GC and while only flopper, phiEcoM_GJ1 and PM1 code for a single tRNA, all of them have exclusively unidirectional coding sequences (CDSs) and code for RNA polymerases, characteristics of the *Autographivirinae* of the *Podoviridae* [64]. However, the verified morphology of ST32, phiEcoM_GJ1, PM1 and PP101, icosahedral head, neck and a contractile tail with tail fibres, classifies them as myoviruses [59,60,62,63]. Based on NT similarity and genome synteny, these seven phages belong to the same, not yet classified, peculiar lineage first described by Jamalludeen *et al*., (2008) [60].

### 3.4. Five novel Microviridae phages species

The singleton lilleven (6.1 kb, 44.4% GC, no tRNA) and the five (four novel) *cluster II* phages (5.3-5.5 kb, 46.9-47.2% GC, no tRNAs) are all *Microviridae*, a family of small ssDNA non-enveloped icosahedral phages (Table 2). Lilleven is closely related to (93.9%, BLAST), have pronounced gene pattern synteny and high AA similarities (89-90%), with the Alphatremicrovirus Enterobacteria phage St1 (Figure S2), and is a novel species of the genus *Alphatremicrovirus*, subfamily *Bullavirinae*. The *cluster II* phages comprise four distinct species, only differing by single nucleotide polymorphisms and in non-coding regions (Figure S2). They have high intra-Gegenees score (88-92%) and share genomic organisation and extensive NT similarity (92.6-94.5-%, BLAST) with the unclassified microvirus Escherichia phage SECphi17 (LT960607), primarily differing by single nucleotide polymorphisms and in noncoding regions (Figure S2, Table 2). *Cluster II* are only related to one recognised phage species *Escherichia virus ID52* (63%, BLAST), genus *Gequatrovirus*, subfamily *Bullavirinae*, with whom they predominantly differ in the region in and around the major spike protein (*gpG*), a distinctive marker of the subfamily *Bullavirinae* involved in host attachment. The phylogenetic analysis separates *cluster II* from the gequatroviruses, and both the Gegenees scores (<22%) and NT similarities (<65% BLAST) are moderately low (Figure S2). Nonetheless, *cluster II* are still, based on pronounced genome organisation synteny and a conserved AA similarity (62-64%, Gegenees) considered to be gequatroviruses (Figure S2).

The sequencing of the microviruses is peculiar, as library preparation with the Nextera^®^ XT DNA kit applies transposons targeting dsDNA. However, during microvirus infection the host polymerase converts the viral ssDNA into an intermediate state of covalently closed dsDNA, which is then replicated in rolling circle by viral replication proteins transcribed by the host RNA polymerase [65]. This intermediate state may have enabled the library preparation. The presence of host DNA in the sequence results indicates an insufficient initial DNase1 treatment, which can be attributed to chemical inhibition or inactivation of the enzyme by adhesion to the sides of wells. Hence, it is reasonable to assume that the extracted microvirus DNA was captured as free dsDNA inside host cells during ongoing infections.

### 3.5. Twenty-five novel Siphoviridae phage species

In spite of similar genome sizes, the large group of *Siphoviridae*, is the most diverse (28 unique, 24 novel) in this study, with GC contents ranging from 43.9-54-6% (Figure 3, Table 2). The majority, *clusters IV*-*V* and singleton jahat, are of the subfamily *Tunavirinae*, while the remaining are allocated into at least three divergent genera, *Dhillonvirus, Jerseyvirus* and *seuratvirus*.

*Cluster IV* (46.7-51.6 kb, 43.9-44.1% GC, no tRNAs), *V* (49.8-51.6 kb, 44.6-44.8% GC, no tRNAs) and jahat (51.1 kb, 45.7% GC, no tRNAs) have low inter-Gegenees scores (7-20%) (Figure 1), yet they form a monophyletic clade and are all closely related (85-94%, BLAST) to published *Tunavirinae*. The *Cluster IV* phages are notable, as they represent a significant increase to the small genus *Hanrivervirus* comprising only the type species *Shigella virus pSf-1* (51.8 kb, 44% GC, no tRNAs) isolated from the Han River in Korea [66]. The common ancestry of *Cluster IV* and pSf-1 is evident by comparable genome sizes, GC contents, genomic organization and substantial NT (86-90%, BLAST) and AA (77-85%, Gegenees) similarities (Table 2, Figure S3). During their differentiation, many deletions and insertions of small hypothetical genes have occurred, most notable is a unique version of a putative tail-spike protein in seven of the *cluster IV* hanriverviruses, indicating divergent host ranges (Figure S3). All the hanriverviruses code for (putative) *dam* and Psf-1 is resistant towards at least six restriction endonucleases [66], suggesting they employ DNA methylation as a defence strategy.

The five novel dhillonviruses of *Cluster XII* (44.4-46.0 kb, 54.2-54.6% GC, no tRNAs) have substantial NT similarities with a group of unclassified dhillonviruses (80-89%, BLAST) (Table 2) and the type species *Escherichia virus HK578* (77-80%, BLAST). As with the hanriverviruses, and as observed by Korf *et al*., (2019) [22], their genomes mainly differ in minor hypothetical genes and in putative tail-tip proteins, indicating divergent host ranges (Figure S4). Based on NT similarity and the presence of the canonical 7-deazaguanine operon the singleton skure (59.4 kb, 44.6% GC, no tRNAs) is a seuratvirus, while the singleton Buks (40.3 kb, 49.7% GC, no tRNAs) is assigned to the genus *Jerseyvirus*, subfamily *Guernseyvirinae*.

#### 3.5.1 Escherichia phage halfdan, a novel lineage within the *Siphoviridae*

Interestingly, the singleton halfdan (42.8 kb, 53.7% GC, no tRNAs) has only miniscule similarity with published phages (12-29%, BLAST). These entail two *Pseudomonas* phages vB_PaeS_SCUT-S3 (MK165657) and Ab26 (HG962376) [67] both *Septimatreviruses*, two *Acinetobacter* phages of the *Lokivirus* IMEAB3 (KF811200) and type species *Acinetobacter virus Loki* [68], and to a lesser degree the unclassified *Achromobacter* phage phiAxp-1 (KP313532) [69]. They have a common genomic organization, yet their intra-Gegenees score is ≤1% and NT similarity is negligent in roughly a third of halfdan’s 57 CDSs (Figure 4b, d). The phylogeny and AA similarities (Gegenees) also indicate a distant relation, although grouping halfdan closer with the lokiviruses (40-43%) than the septimatreviruses (33-34%) (Figure 4a, c). The genome of halfdan is mosaic, resembling the lokiviruses in the structural region and the septimatreviruses in the replication region (Figure 4d). Remarkably, halfdan has no NT similarity with known *Escherichia* phages, which could be an indication of *E. coli* not being the natural host, and that halfdan may have to transcended species barriers. However, host range, morphology and other physical characteristics are beyond the scope of this study, and will be determined in future lab-based studies. Notably, both halfdan, Loki and Ab26 code for a (putative) MazG, although there is negligible sequence similarity (Figure 4d). Phage encoded MazG is hypothesized to be involved in restoring protein synthesis in a starved cell, in order to keep the cell alive, and ensure optimal replication conditions [70]. The phylogeny, whole genome alignments and low NT and AA similarities suggests that halfdan is distinct from other known phages (Figure 4). Accordingly, as per the ICTV guidelines, halfdan is the first phage sequenced of a novel *Siphoviridae* genus. Yet, halfdan is also, by its mosaicism, an indicator of the genetic continuum of phages questioning the validity of taxonomic interpretations.

**Figure 4.**
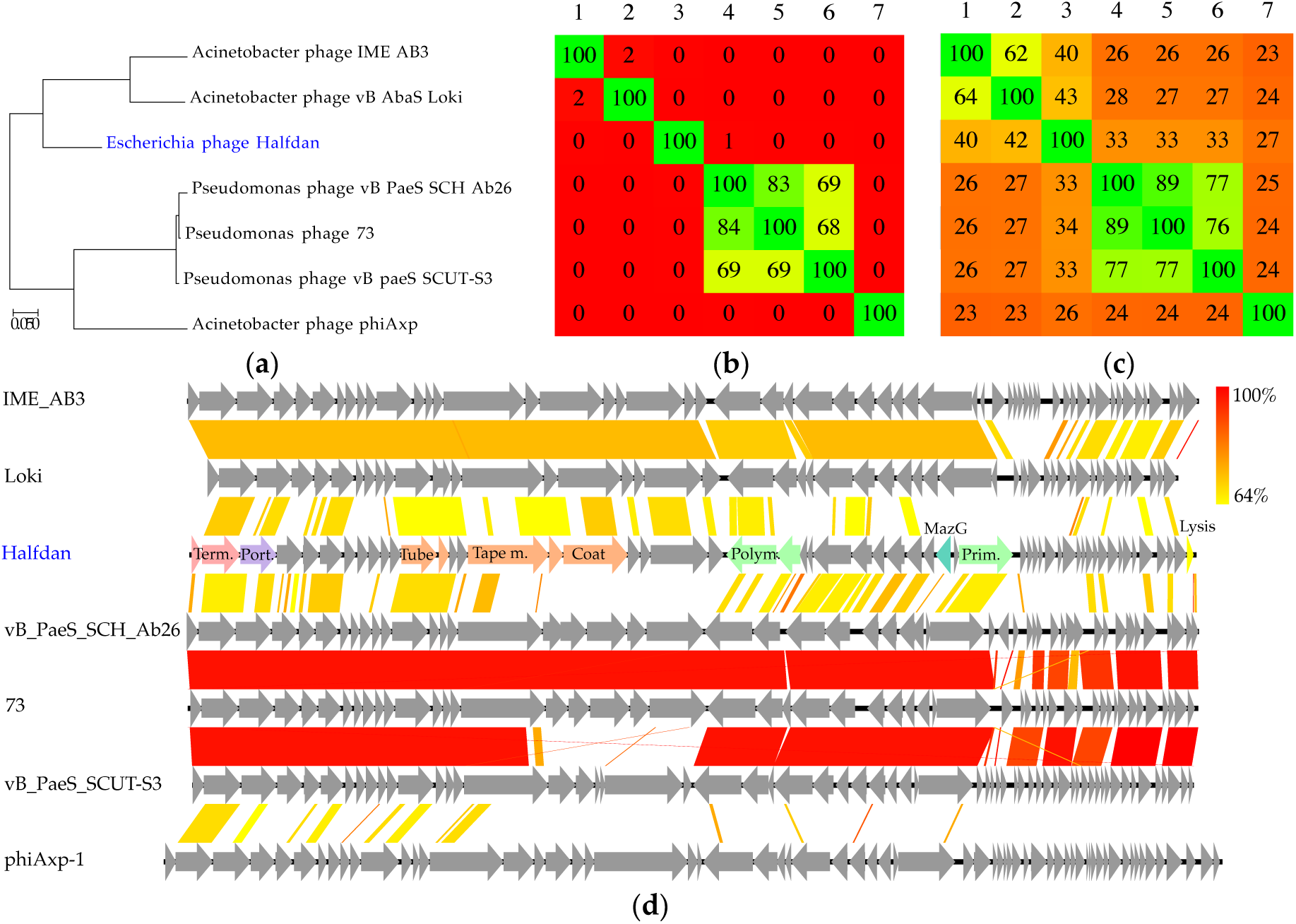
(**a**) Phylogenetic tree (Maximum log Likelihood: −7678.71, large terminase subunit), scalebar: substitutions per site. (**b**) Phylogenomic nucleotide distances (Gegenees, BLASTn: fragment size: 200, step size: 100, threshold: 0%). (**c**) Phylogenomic amino acid distances (Gegenees, tBLASTx: fragment size: 200, and step size: 100, threshold: 0%). (**d**) Pairwise alignment of phage genomes (Easyfig, BlASTn), the colour bars between genomes indicate percent pairwise similarity as illustrated by the colour bar in the upper right corner (Easyfig, BlASTn). Gene annotations are provided for phages identified in this study and deposited in GenBank.

### 3.6. Nine novel Podoviridae phage species

The nine novel *Podoviridae* (c*luster XIV*, 40.7-42.5 kb, 55.4-55.7% GC, no tRNAs), all with the hallmarks of the *Autographvirinae i.e*. unidirectionally encoded genes and RNA polymerases [64], have conserved genome organisation with the type species of the *Phikmvvirus, Pseudomonas virus phiKMV* (42.5 kb, 62.3% GC, no tRNAs) [71], but almost no NT similarity (<1%, BLAST). Besides the unclassified *Autographvirinae* Enterobacteria phage J8-65 (NC_025445) (40.9 kb, 55.7% GC, no tRNAs), with which *Cluster XIV* has considerable NT similarity (69-93%, BLAST) and similar GC content, *Cluster XIV* only share >5% nucleotide similarity (40-42%, BLAST) with the *Phikmvvirus Pantoea virus Limezero* (43.0 kb, 55.4% GC, no tRNAs) [72] (Figure 5, Table 2). The *Phikmvvirus* consists of four species, *phiKMV, Limezero, Pantoea virus Limelight* (44.5 kb, 54% GC, no tRNAs) [72] and *Pseudomonas virus LKA1* (41.6 kb, 60.9% GC, no tRNAs) [73].

**Figure 5.**
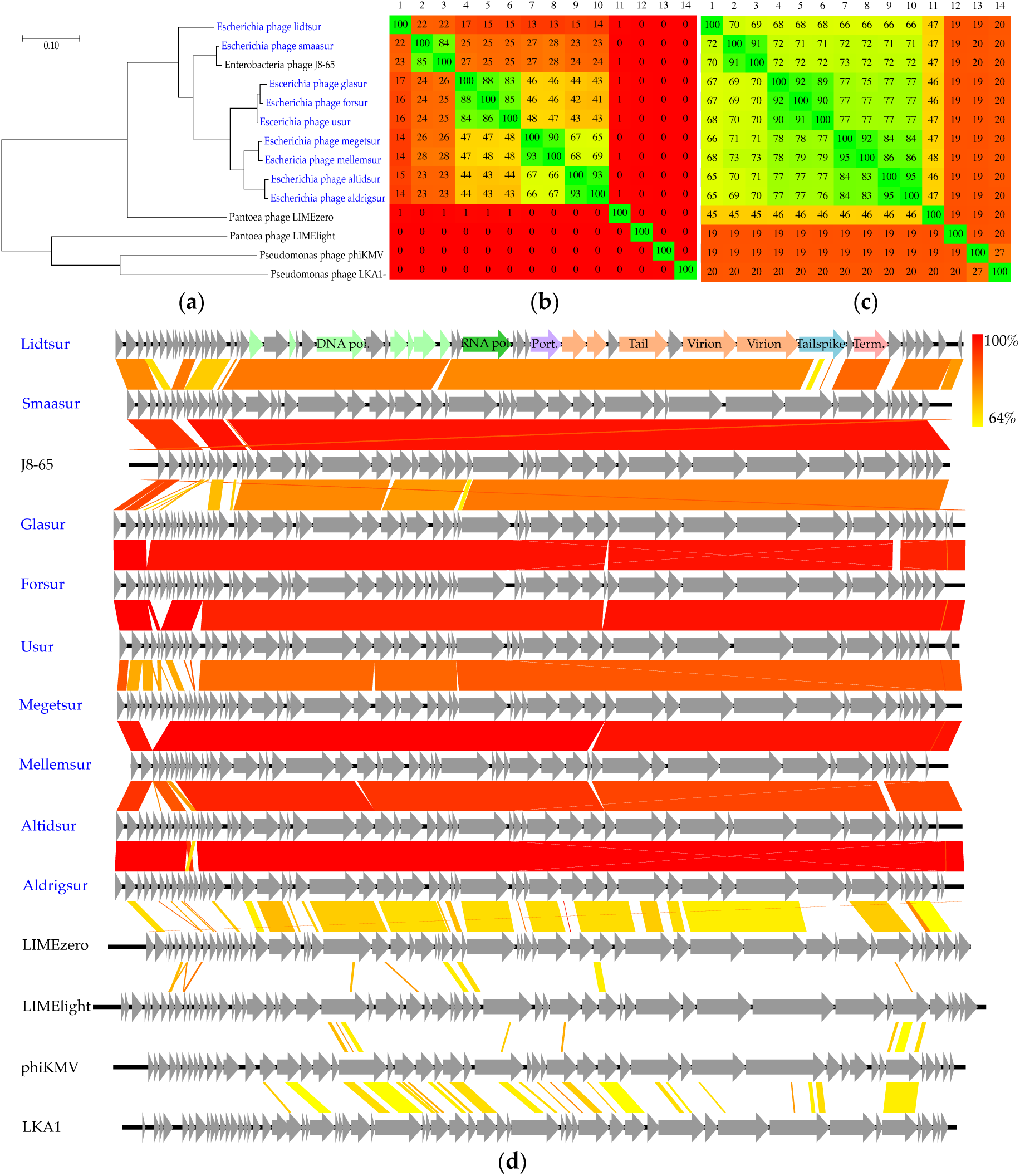
(**a**) Phylogenetic tree (Maximum log Likelihood: −11728.26, large terminase subunit), scalebar: substitutions per site. (**b**) Phylogenomic nucleotide distances (Gegenees, BLASTn: fragment size: 200, step size: 100, threshold: 0%). (**c**) Phylogenomic amino acid distances (Gegenees, tBLASTx: fragment size: 200, and step size: 100, threshold: 0%). (**d**) Pairwise alignment of phage genomes (Easyfig, BLASTn), the colour bars between genomes indicate percent pairwise similarity as illustrated by the colour bar in the upper right corner (Easyfig, BlASTn). Gene annotations are provided for phages identified in this study and deposited in GenBank.

*Cluster XIV* and J8-65 form a diverse monophyletic clade, with a substantial amount of deletions and insertions between them, subdividing into three sub-clusters with intra-Gegenees scores ≤28% (Figure 5a, d). Phage lidtsur is singled-out and also codes for a unique version of tailspike colanidase, smaasur resembles J8-65 phage and the rest group together (Figure 5). Still, the three sub-clusters have for a large part conserved AA sequences (66-72%, Gegenees) (Figure 5b, c, d). Interestingly, this also applies to *Limezero*, with whom they have a Gegenees NT score ≤1%, but an AA similarity of 45-48%, supporting the phylogenetic grouping of *Limezero* and *cluster XIV* (Figure 5a, b, c). Based on phylogeny, limited NT and low AA similarities it is evident that there is a very distant relation between *cluster XIV* (and J8-65) and the phikmvviruses (Figure 5). Hence, *cluster XIV* (and J8-65) are not immediately, according to the ICTV guidelines, considered phikmvviruses. However, the *Phikmvvirus* already includes phages with minuscule DNA homology infecting divergent hosts [72,73]. The genus delimitation is based on overall genome architecture with the location of a single-subunit RNA polymerase gene adjacent to the structural genes, and not in the early region as in T7-like phages [16]. Another characteristic feature of the *phikmvvirus* is the presence of direct terminal repeats, though not yet verified in all members [72]. Read abundance in non-coding regions in genomic ends of *cluster XIV* genomes suggests they also have direct terminal repeats of a few hundred bp. In conclusion, we consider the *cluster XIV* phages (and J8-65) to be of the same lineage as the phikmvviruses, but contemplate that this genus may at some point be divided into at least two independent genera, as we learn more of the nature of the genes which distinguish *cluster XIV* from the other phikmvviruses on both NT and AA level and account for more 50% of their genomes.

### 3.7. Unclassified bacterial viruses represent two novel lineages

The four novel phages sortsyn of *cluster XIII* and the three singletons sortsne, sortkaff (41.9-42.5 kb, 59-60% GC, no tRNAs) and skarpretter (42.0 kb, 55.8% GC, no tRNAs) all have small genomes and high GC contents (Figure 3) and are suggested to represent two distinct novel lineages. They only share considerable (≥6%, BLAST) NT similarity with four phages, Enterobacteria phage IME_EC2 (KF591601)(41.5 kb, 59.2% GC, no tRNAs) isolated from hospital sewage [74], Salmonella phage lumpael (MK125141)(41.4 kb, 59.5% GC, no tRNAs) isolated from wastewater of the same sample-set in a previous study, Klebsiella phage vB_KpnS_IME279 (MF614100) (42.5 kb, 59.3% GC, no tRNAs) and Escherichia phage C130_2 (MH363708)(41.7 kb, 55.4% GC, no tRNAs) isolated from cheese [75].

Curiously, IME_EC2 is a confirmed (TEM) *Podoviridae*, with a short non-contractile tail [74], while C130_2 is a confirmed (TEM) *Myoviridae* with a long contractile tail [75]. Lumpael and vB_KpnS_IME279 are uncharacterised, yet both are assigned as *Podoviridae* in GenBank [17]. Although sortsne, sortkaff, sortsyn, vB_KpnS_IME279, IME_EC2 and lumpael form a monophyletic clade, the Gegenees score (5-15%) between sub-clusters is surprisingly low (Figure 6a, b). Still, these six phages have comparable genome sizes (41.5-42.5 kb) and organisation, including similar relatively small-sized structural proteins, *dam* and *dcm* genes, equally high GC contents (59.5±0.5%) and relatively high AA similarities (49-91%) (Figure 6a, c, Table 2). Hence, they are clearly of the same lineage and likely to resemble IME_EC2 in having *Podoviridae* morphology. This group is also clearly distinct from all other known phages (<5%, BLAST) and as such constitute a novel genus, with a delimitation to be determined by future physical characterisation. Skarpretter and C130_2 form a monophyletic clade and both have slightly, though not significantly (*P* = 0.38), lower GC contents (55.6±0.2%), indicating a common ancestry and host. The conserved genome organisation and clustering of skarpretter with the other *Podoviridae* (Figure 1, 6d), suggests skarpretter also has *Podoviridae* morphology. Still, its closest relative C130_2 is described as a *Myoviridae* [75], leaving the true morphology of skarpretter a conundrum only resolved by TEM imaging. According to ICTV guidelines both skarpretter and C130_2 are representatives of novel linages, as they are both clearly distinct from all other known phages. Skarpretter and C130_2 have very low NT (1% Gegenees, 38% BLAST) and AA similarities (43%) (Figure 6b, c). Indeed, skarpretter has ≤20% NT similarity (BLAST) with all other published phages. In addition, skarpretter codes for a putative *hflc* gene possibly inhibiting proteolysis of essential phage proteins, located in the replication region (Figure 6d). This gene, has to our knowledge never before been observed in phages, but occurs in almost all proteobacteria, including *E. coli* MG1655 [76].

**Figure 6.**
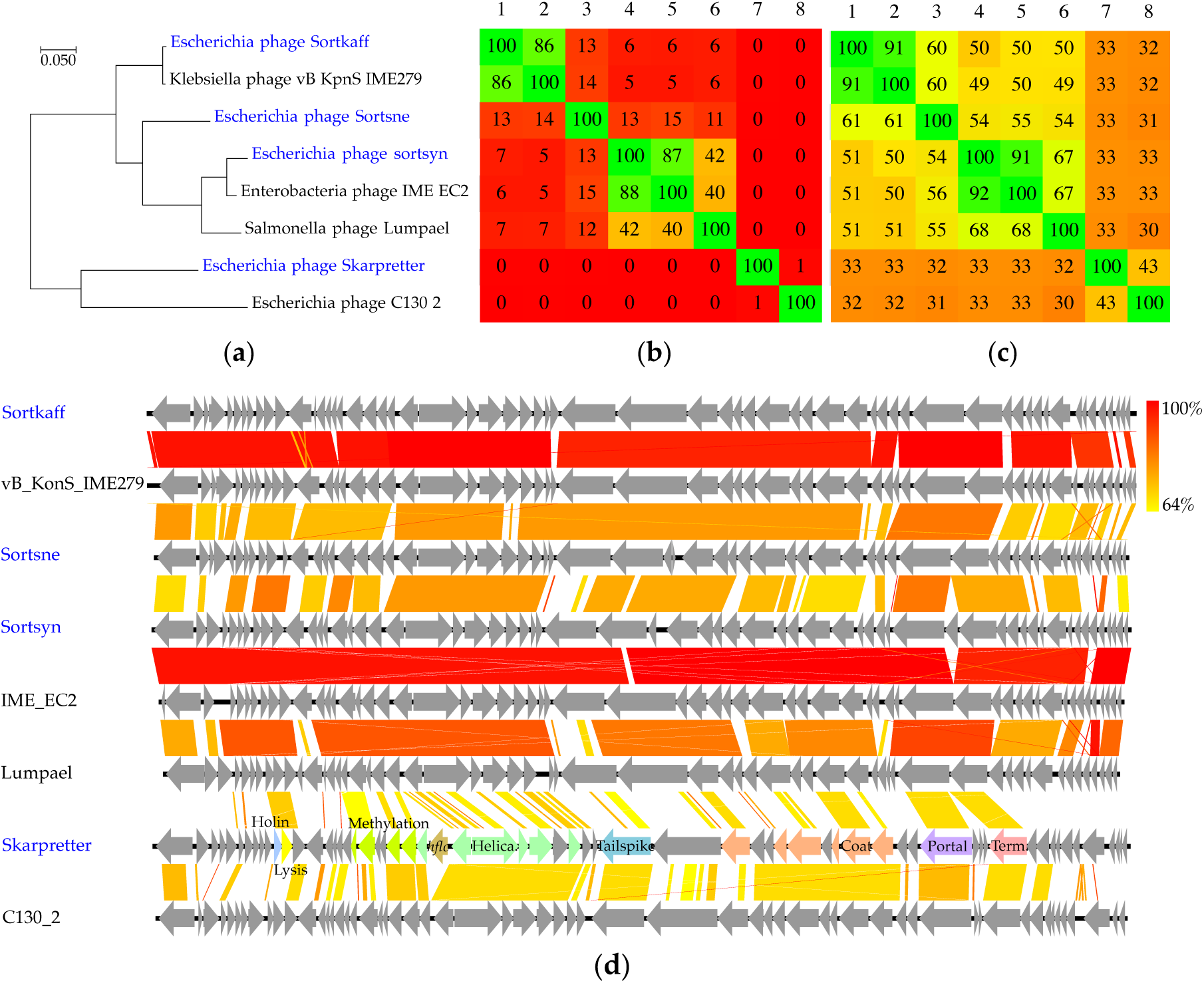
(**a**) Phylogenetic tree (Maximum log Likelihood: −8023.43, large terminase subunit), scalebar: substitutions per site. (**b**) Phylogenomic nucleotide distances (Gegenees, BLASTn: fragment size: 200, step size: 100, threshold: 0%). (**c**) Phylogenomic amino acid distances (Gegenees, tBLASTx: fragment size: 200, and step size: 100, threshold: 0%). (**d**) Pairwise alignment of phage genomes (Easyfig, BlASTn), the colour bars between genomes indicate percent pairwise similarity as illustrated by the colour bar in the upper right corner (Easyfig, BlASTn). Gene annotations are provided for phages identified in this study and deposited in GenBank.

## 4. Conclusion

By screening Danish wastewater, we identified no less than 104 unique *Escherichia* MG1655 phages, but predict the species richness to be at least in the range of 160-420, and even higher if including phages infecting other *Escherichia* species, though it is expected to fluctuate drastically over time and both within and between treatment facilities. Among the unique phages 91 represent novel phage species of at least four different families *Myoviridae, Siphoviridae, Podoviridae* and *Microviridae*. The diversity of these phages is striking, they vary greatly in genome size and have a very broad GC-content range - possibly prompted by former or current alternate hosts. These findings add to our growing understanding of phage ecology and diversity, and through classification of these many phages we come yet another step closer to a more refined taxonomic understanding of phages. Furthermore, the numerous and diverse phages isolated in this study, all lytic to the same single strain, serve as an excellent opportunity to learn important phage-host interactions in future studies. These include, but are not limited to lysogen induced phage immunity, host-range and anti-RE systems. Finally, apart from substantial contributions to known genera, we came upon several unclassified lineages, some completely novel, which in our opinion constitute novel phage genera. We consider sortregn, sortkaff, sortsyn and sortsne together with lumpael, vB_kpnS_IME278 and IME_EC2 to constitute a novel genus within the *Podoviridae*. The *Myoviridae* flopper joins an interesting group of not yet classified phages with *Myoviridae* morphology and *Autographvirinae* characteristics, just as *cluster VII* together with five unclassified phages represent a novel unclassified genus within the *Myoviridae*. Lastly, we consider the *Siphoviridae* halfdan and the unclassified bacterial virus skarpretter to be the first sequenced representatives of each their novel genus. In conclusion, this study shows that uncharted territory still remains for even well-studied phage-host couples.

## Supplementary Materials

Figure S1: Microviridae, Figure S2: Hanriverviruses, Figure S3: Dhillonviruses, Table 1 Auxiliary metabolism genes.

## Acknowledgments

This study had not been possible without the much-appreciated contribution by the many members of the Danish Water and Wastewater Association (DANVA), who kindly supplied us with time-series of wastewater samples from their treatment facilities. A special thanks to the Billund and Grindsted treatment facilities of Billund Vand & Energi, Lynetten, Avedøre and Damhusåen treatment facilities of BIOFOS, Kolding treatment facility of BlueKolding, Esbjerg Øst, Esbjerg Vest, Ribe, Varde og Skovlund treatment facilities of DIN Forsyning, Drøsbro, Hadsten, Hammel, Hinnerup and Voldum treatment facilities of Favrskov Forsyning, Haerning treatment facility of Herning Vand, Hillerød treatment facility of Hillerød Forsyning,Lemvig treatment facility of Lemvig Vand og Spildevand, Bevtoft, Gram, Haderslev, Halk, Jegerup, Nustrup, Over Jerstal, Skovby, Skrydstrup, Sommersted, Vojens and årøsund treatment facilities of Provas, Marselisborg, Egå and Viby treatment facilities of Aarhus vand, Hedensted, Juelsminde and Tørring treatment facilities of Hedensted Spildevand, Ejby Mølle, Bogense, Hofmansgave, Hårslev, Nordvest, Nordøst, Otterup and Søndersø treatment facilities of VandCenterSyd, Øst and Vest treatment facilities of Aalborg Forsyning and finally Helsingør treatment facility of Forsyning Helsingør.

## SUPPLEMENTARY FIGURES AND TABLES

**Figure S1.**
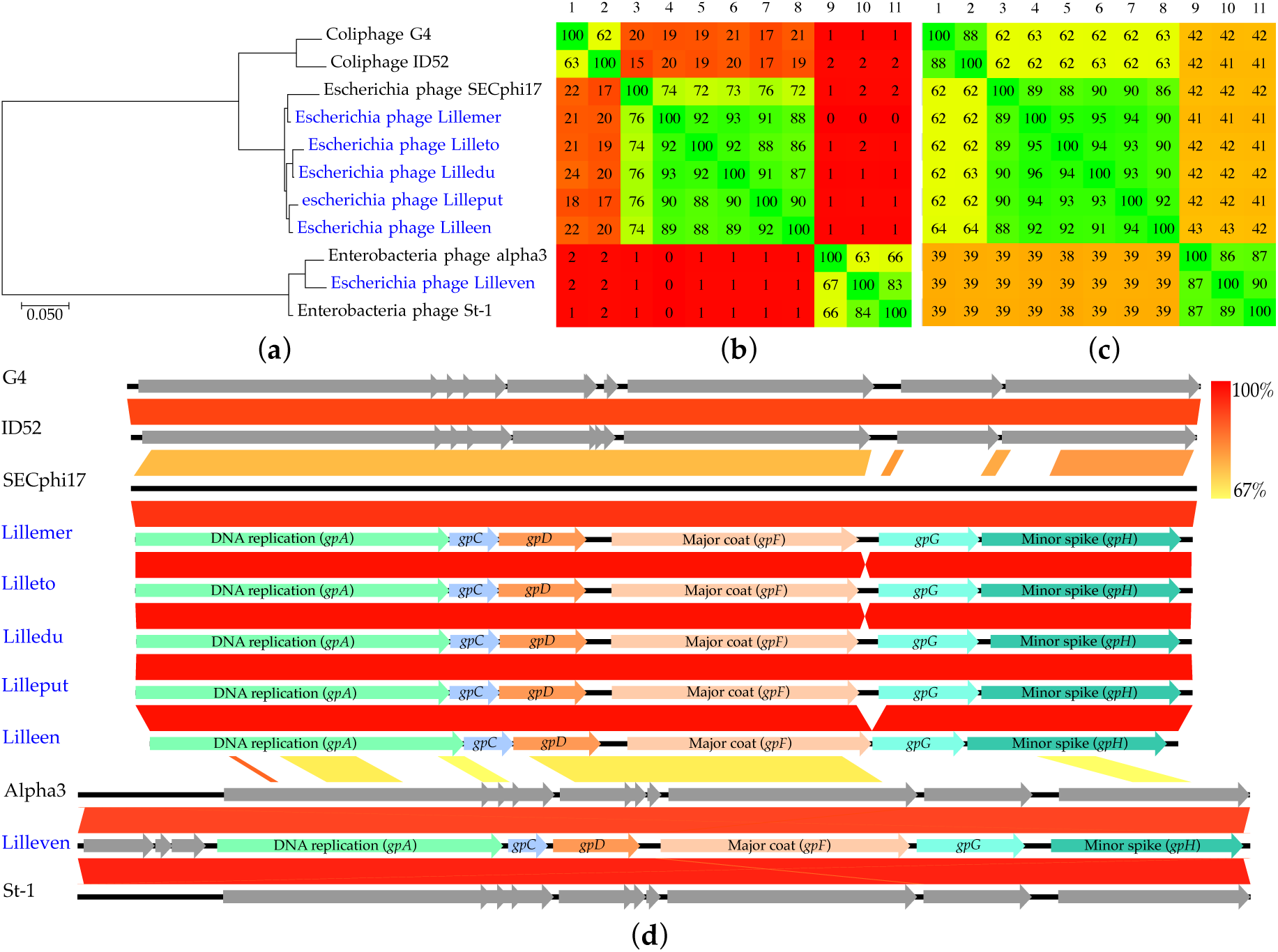
Microviridae. **Figure S4** (a) Phylogenetic tree (Maximum log Likelihood: −6065.3, DNA replication protein gpA), scalebar: substitutions per site (b) Phylogenomic nucleotide distances (Gegenees, BLASTn: fragment size: 200, step size: 100, threshold: 0%). (c) Phylogenomic amino acid distances (Gegenees, tBLASTx: fragment size: 200, and step size: 100, threshold: 0%). (d) Pairwise alignment of phage genomes, the colour bars between genomes indicate percent pairwise similarity as illustrated by the colour bar in the upper right corner (Easyfig, BlASTn). Gene annotations are provided for phages identified in this study and deposited in GenBank.

**Figure S2.**
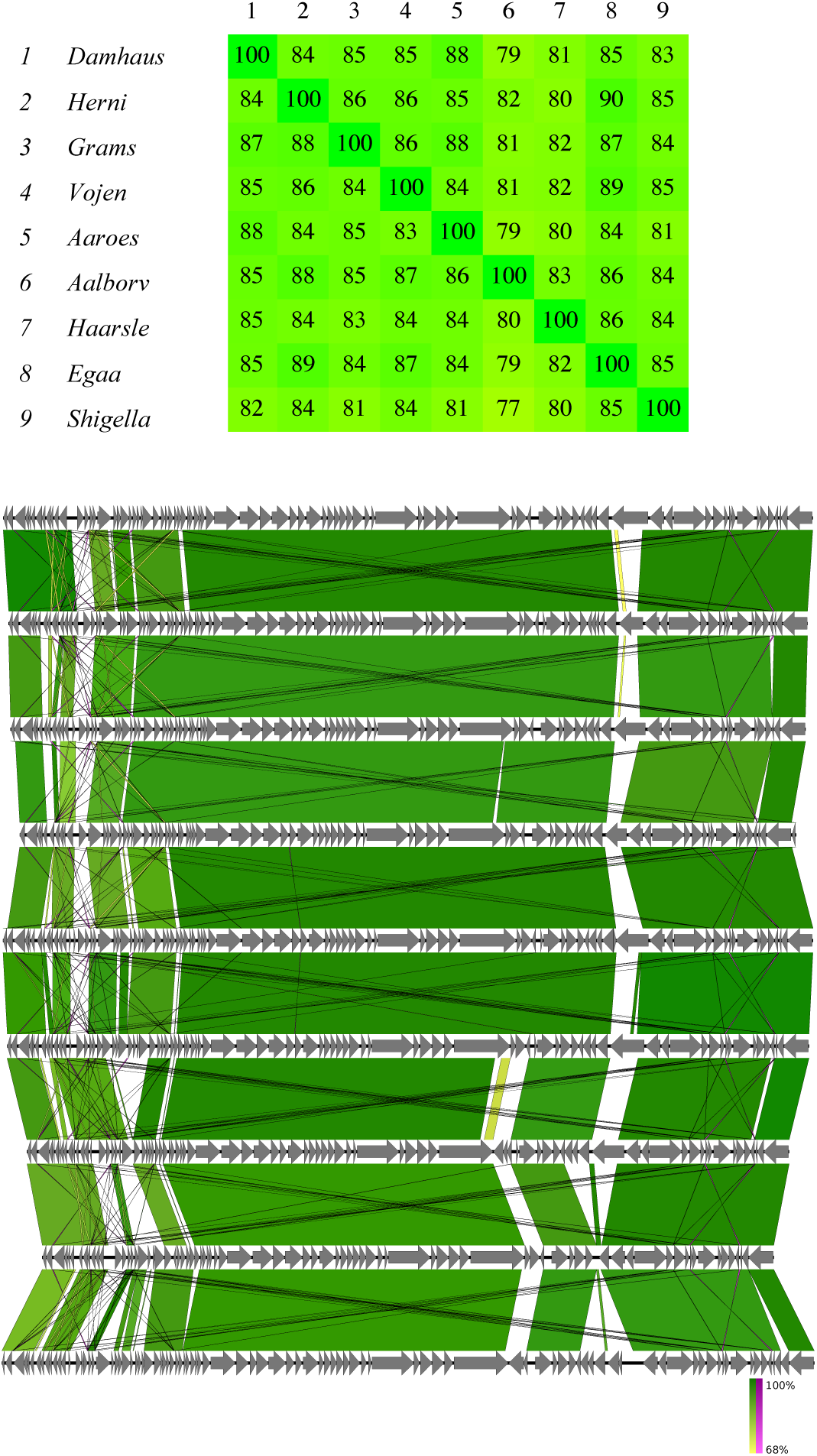
Hanrivervirus. Phylogenomic amino acid distances (Gegenees, tBLASTx: fragment size: 200, and step size: 100, threshold: 0%). and Pairwise alignment of phage genomes (Easyfig, BlASTn), the colour bars between genomes indicate percent pairwise similarity as illustrated by the colour bar in the lower right corner (Easyfig, BlASTn). Genome order is the same in both analyses.

**Figure S3.**
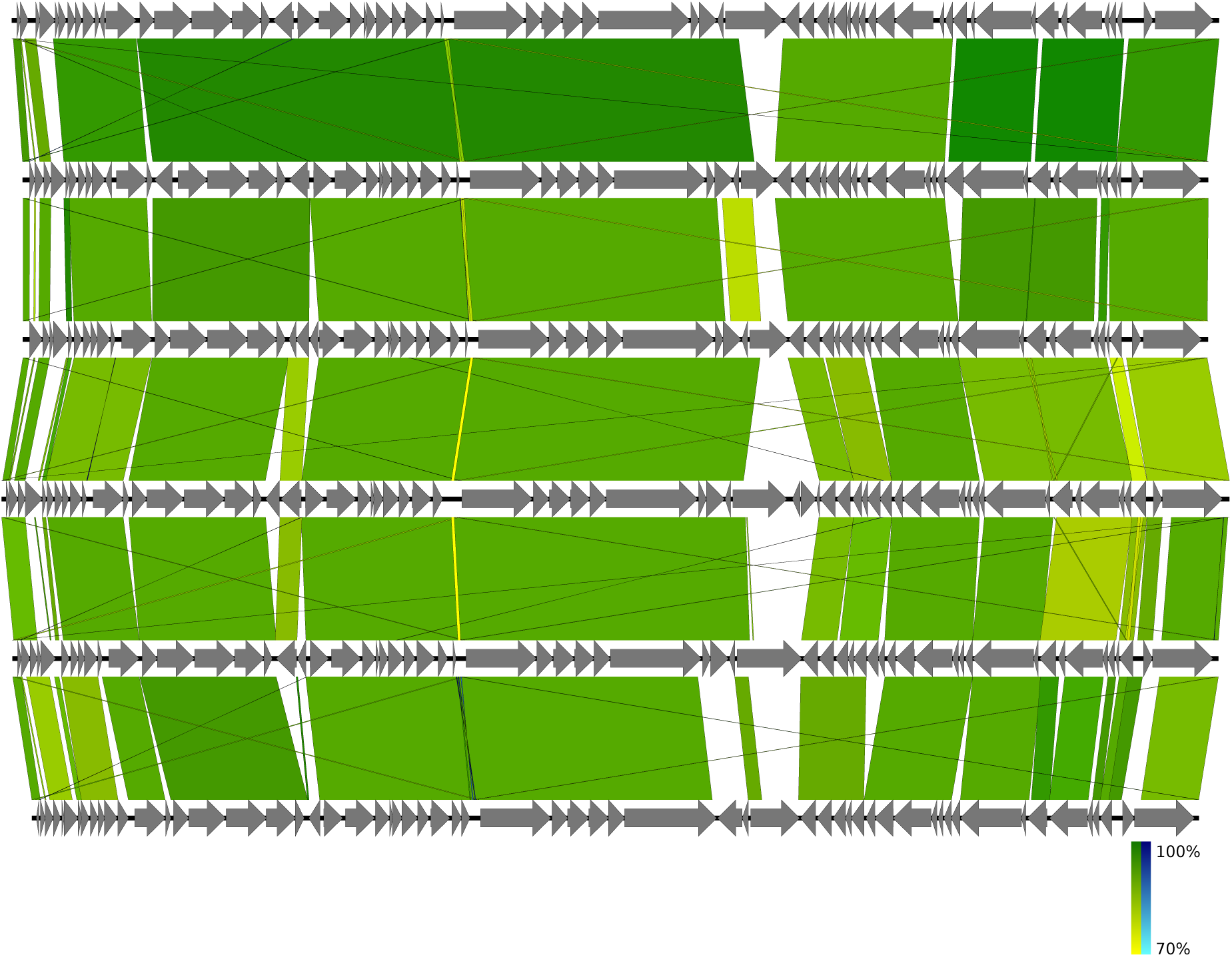
Dhillonvirus. Pairwise alignment of phage genomes (Easyfig, BlASTn), the colour bars between genomes indicate percent pairwise similarity as illustrated by the colour bar in the lower right corner (Easyfig, BlASTn).

**Table S1.**
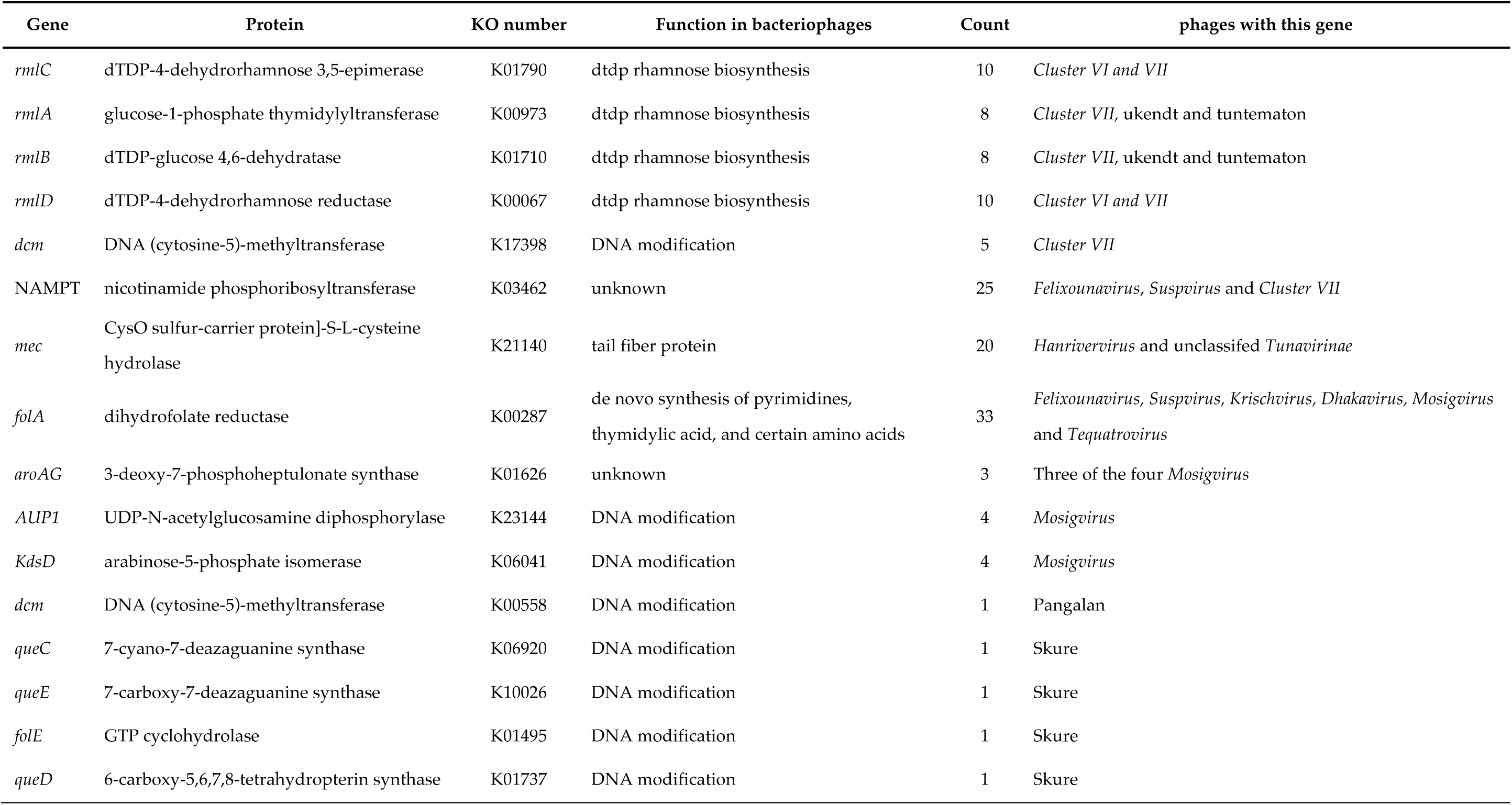
Auxiliary metabolism genes.

